# Reclassification of probiotic *Lactobacillus acidophilus* NCIMB 30184 as *Lactobacillus helveticus* and *Lactobacillus casei* NCIMB 30185 as *Lacticaseibacillus paracasei*

**DOI:** 10.1101/2022.10.17.512536

**Authors:** A. Rodenes, B. Casinos, B. Alvarez, J.F. Martinez-Blanch, V. Vijayakumar, G. Superti, R. Day, E. Chenoll

## Abstract

The strains NCIMB 30184 and NCIMB 30185, originally designated as *Lactobacillus acidophilus and Lactobacillus casei* respectively, have been studied in published pre-clinical and clinical studies. Their nucleotide sequences of the 16S rRNA were compared with public databases and were preliminarily found to have the highest similarity with *Lactobacillus helveticus* and *Lacticaseibacillus paracasei* species respectively. Subsequently, total 16S and 23S rRNA genes were compared with their most similar taxa, and the preliminary identification as *Lactobacillus helveticus* and *Lacticaseibacillus paracasei* were confirmed.

## INTRODUCTION

Initially proposed by Beijerinck in 1901 as Gram-positive, homofermentative, thermophilic and non-spore-forming rods, genus *Lactobacillus* has experienced multiple taxonomical reassignments. Initially based on phenotypic traits [1], its taxonomic assignment criteria has been modified throughout the last decades mostly based on molecular approaches. Sequencing of 16S rRNA genes has been established in bacterial taxonomy as a marker gene in phylogenetic studies for classification and nomenclature purposes [2]. However, *Lactobacillus* is a very complex genus, in which the initial phenotypic differentiations are highly variable within clades [3,4] with an ever increasing number of new species and re-designations [5,6]. This heterogeneity was finally studied as a whole by Zheng and coworkers [7], who divided the genus in 23 different genera. All of these reassignments and changes have made new taxonomical evaluation of the strains necessary and have reinforced the taxonomical complexity of this genus in its identification.

Strain NCIMB 30184, originally designated as lactic acid bacteria *Lactobacillus acidophilus*, has been studied in published pre-clinical and clinical studies [8,9]. An isolate from human origin, probiotic strain PXN35 was deposited at National Collection of Industrial, Food and Marine Bacteria (NCIMB, Aberdeen, United Kingdom) on the 7^th^ of April 2006 as *L. acidophilus*.

Strain NCIMB 30185 was identified initially as *Lactobacillus casei* and its functionality was informed in [8,9]. The probiotic strain PXN37 was deposited at NCIMB collection on the 7^th^ of April 2006 as *L. casei*.

In both NCIMB 30184 and NCIMB 30185 strains, *in vitro* microbiological and molecular evaluation performed showed that the genetic characteristics presented by the isolates differed from their original designation. In order to determine the correct identification, the strains were revived from the original master cell banks and 16s rRNA and 23s rRNA gene sequencing analysis were performed to allow comparison with closely related species and determine the correct classification of these probiotic strains.

## MATERIALS AND METHODS

### Culture and growth conditions

The long-term storage stock of the strains from NCIMB (vial dated 22nd October 2009 for PXN 35 and 04^th^ November 2002 for PXN 37)) were revived by culturing in MRS (Oxoid) at 30 °C under microaerophilic conditions. Working cell banks were conserved in 20 % (v/v) glycerol at −80 °C. Microbiological evaluation was applied by standard methodology.

### Nucleotide sequence analysis

DNA from pure culture was extracted following the guanidium thiocyanate method isolation [10], spectrophotometrically quantified (Nanodrop) and adjusted to the final concentration with ultra-pure water (Sigma-Aldrich, Madrid, Spain). 16s rRNA gene sequencing was run following Chenoll et al [11]. Almost full-length 16S rRNA gene fragment was amplified from genomic DNA by using the following primers: 616 Valt (5′-AGA GTT TGA TYMTGG CTC AG-3′) and 630R (5′-CAK AAA GGA GGT GATCC-3′). PCR amplifications were conducted in a solution containing 1× PCR buffer, 100 μmol/l of each dNTP, 1 μmol/l of each primer, 1.5 mmol/l MgCl_2_, 0.5 U of thermoestable Biotaq™ DNA polymerase (BioLine, Randolph, MA) and 5 μl of DNA template, in a final volume of 50 μl. Amplification conditions were: 5 min at 94 °C, 35 cycles of 30 sec at 94 °C, 60 sec at 56 °C and 90 sec at 72 °C, and a final extension of 10 min at 72 °C. Almost full-length 23S rRNA gene was in vitro amplified using three universal primer sets following Elizaquivel and coworkers [12]: 1027V/504R, and Ibrahim et al. 2001 [13]: 1104F/1930R and 1522F/130R (all amplification primers are listed in Table 1). PCR amplifications were carried as described above. Amplification conditions were 5 min at 94 °C, 35 cycles of 30s at 94°C, 45 s at 52 °C and 45 s (with a 5 s increment in each cycle) at 72 °C for elongation, and a final extension of 10 min at 72 °C. All reactions were carried out in Mastercycler nexus 2x system thermal cycler (Eppendorf). Controls devoid of DNA were simultaneously included in the amplification process. The integrity of the PCR products was checked by single bands detection following electrophoresis for 1 h at 100 V in 1.4% (wt/vol) agarose gels in Tris-borate-EDTA buffer.

**Table 1.**
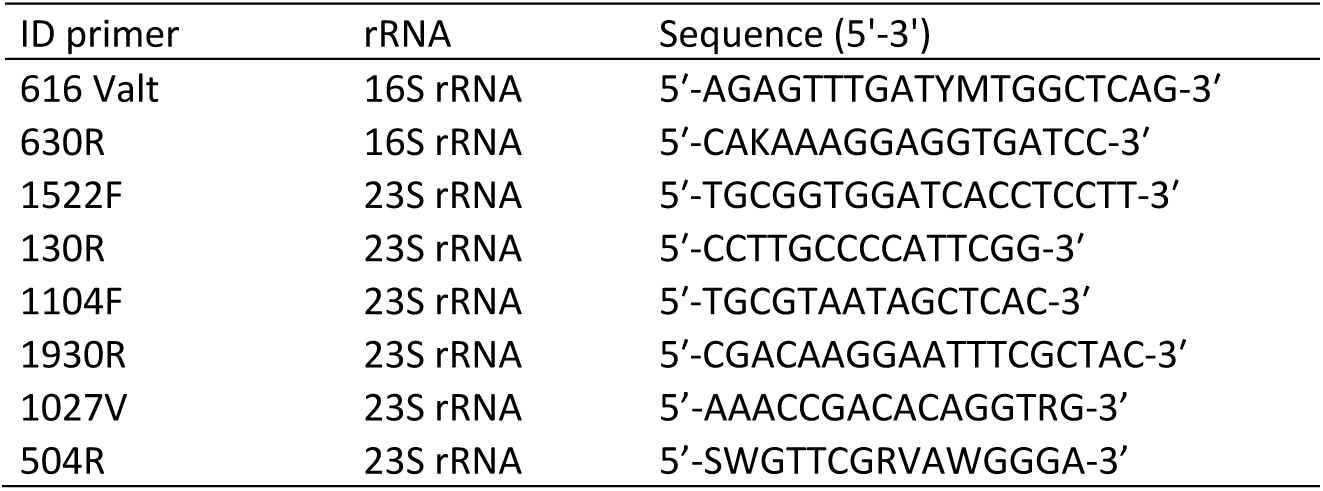
Primers sequences to amplify 16S and 23S rRNA.

### Sequencing of the 16S and 23S rRNA genes and sequence data analysis

Almost full sequences of the 16S and 23S rRNA genes were amplified separately as indicated above. Purification of amplification products was applied by QIAquick PCR purification kit (QIAGEN) and subsequent sequencing reactions were performed using the Big Dye Terminator (version 3.1) cycle sequencing kit (Applied Biosystems) in a premixed format. The 16S rRNA and 23S rRNA sequences determined in this study have been deposited in the GenBank data base (PXN35 accession reference ON495975 and ON495977 respectively, and PXN37 accession reference OP623495 and OP623508 respectively).

The 16S and 23S rRNA sequences were aligned separately and initially inspected by eye and compared by the use of the online tool BLAST (http://blast.ncbi.nlm.nih.gov/Blast.cgi). The strains were tentatively identified on the basis of highest scores. For each strain, two phylogenetic trees were constructed based on 16S and 23 rRNA genes with sequences from type strains (Tables 2-5). Sequences were downloaded from NCBI database and *Bifidobacterium longum* ATCC 15707 was used as outgroup. T-Coffee multiple sequence alignment software [14] was used to align the selected sequences. The jModelTest algorithm [15,16] was used to statistically select the best nucleotide substitution model. In this case, the selected model was TPM2uf+G. PhyML software [17] was then used to check and build the 16S rRNA phylogenetic tree. Finally, the iTOL v5 interactive platform [18] was used for its visualization. Percentage of identity among sequences was obtained by Clustal Omega v2.1 package with default parameters [19].

**Table 2.**
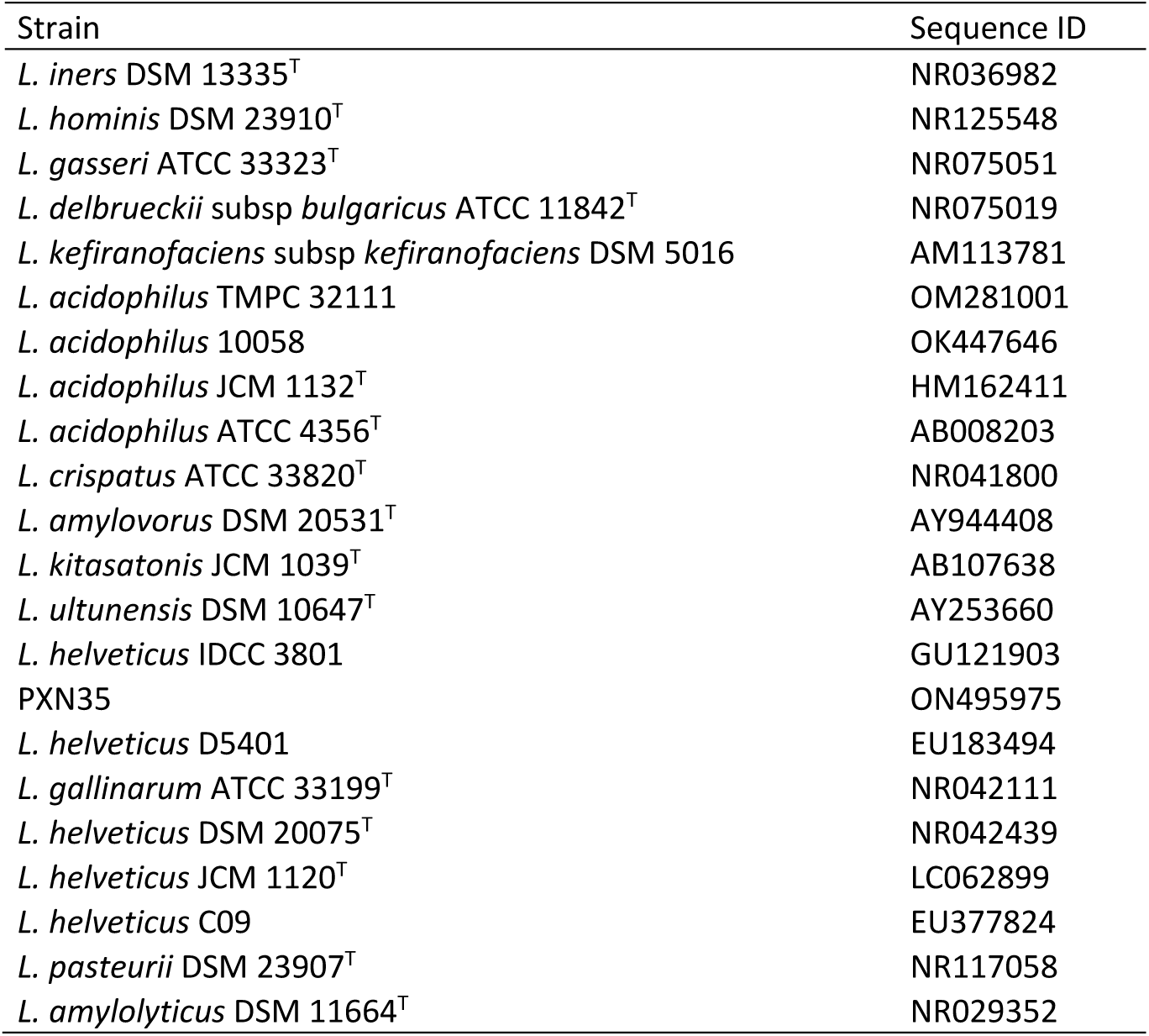
PXN35 16s rRNA gene sequences used for the study.

**Table 3.**
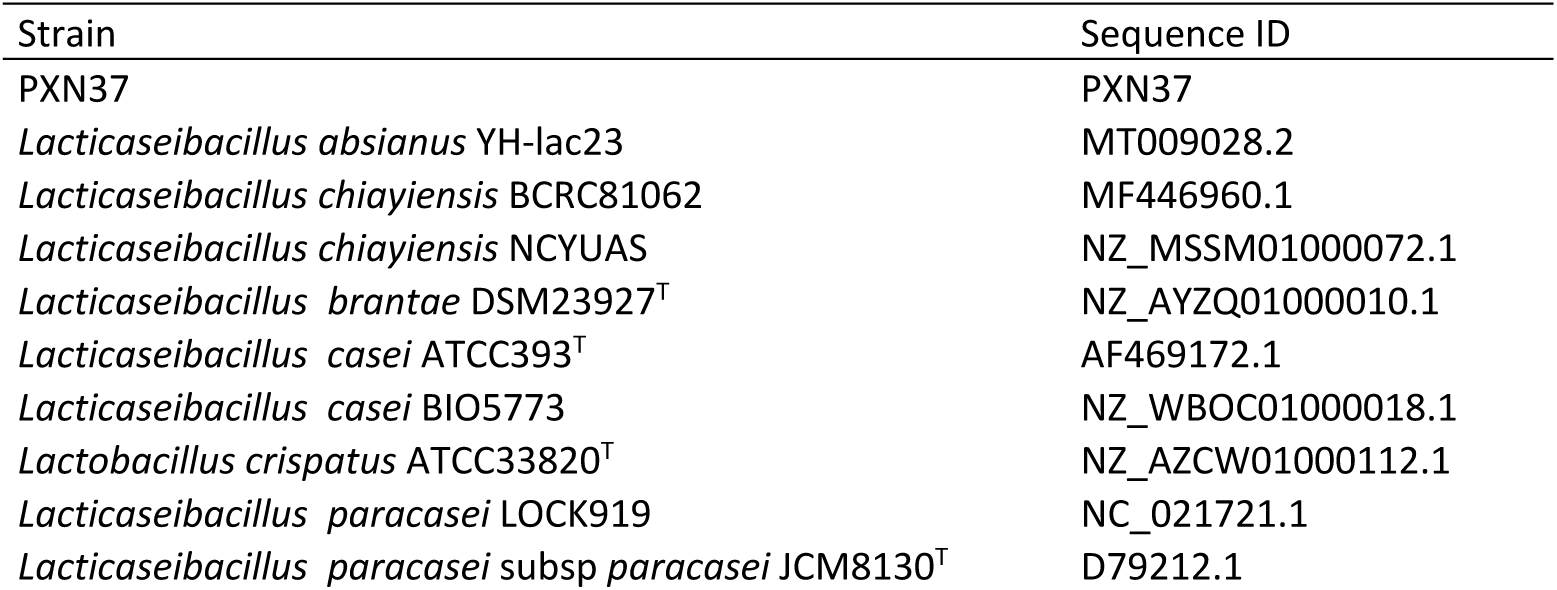
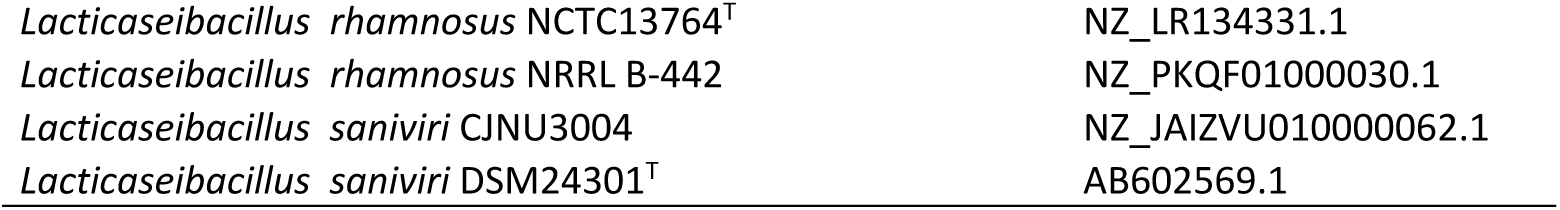
PXN37 16s rRNA gene sequences used for the study.

| Strain | Sequence ID |
| --- | --- |
| PXN37 | PXN37 |
| *Lactocaseibacillus absianus* YH-lac23 | MT009028.2 |
| *Lactocaseibacillus chiayiensis* BCRC81062 | MF446960.1 |
| *Lactocaseibacillus chiayiensis* NCYUAS | NZ_MSSM01000072.1 |
| *Lactocaseibacillus brantae* DSM23927<sup>T</sup> | NZ_AYZQ01000010.1 |
| *Lactocaseibacillus casei* ATCC393<sup>T</sup> | AF469172.1 |
| *Lactocaseibacillus casei* BIO5773 | NZ_WBOC01000018.1 |
| *Lactobacillus crispatus* ATCC33820<sup>T</sup> | NZ_AZCW01000112.1 |
| *Lactocaseibacillus paracasei* LOCK919 | NC_021721.1 |
| *Lactocaseibacillus paracasei* subsp *paracasei* JCM8130<sup>T</sup> | D79212.1 |
| *Lactcaseibacillus rhamnosus* NCTC13764<sup>T</sup> | NZ_LR134331.1 |
| *Lactcaseibacillus rhamnosus* NRRL B-442 | NZ_PKQF01000030.1 |
| *Lactcaseibacillus saniviri* CJNU3004 | NZ_JAIZVU010000062.1 |
| *Lactcaseibacillus saniviri* DSM24301<sup>T</sup> | AB602569.1 |

**Table 4.** PXN35 23S rRNA gene sequences used for the study.

| Strain | Sequence ID |
| --- | --- |
| *L. iners* DSM 13335<sup>T</sup> | NZ_GG700803.1 |
| *L. gasseri* 63 AM | NR_076437.1 |
| *L. hominis* DSM 23910 | NZ_AYZP01000029.1 |
| *L. delbrueckii* subsp *bulgaricus* ATCC 11842<sup>T</sup> | NR_077006.1 |
| *L. amylolyticus* DSM 11664<sup>T</sup> | Y17360.1 |
| *L. pasteurii* DSM 23907<sup>T</sup> | NZ_PUFQ01000021.1 |
| *L. kefiranofaciens* subsp *kefiranofaciens* JCM 6985 | NZ_BAMG01000080.1 |
| *L. ultunensis* Kx293C1<sup>T</sup> | NZ_CP059829.1 |
| *L. acidophilus* NCFM | NR_076355.1 |
| *L. amylovorus* GRL 1112 | NR_076802.1 |
| *L. kitasatonis* JCM1039<sup>T</sup> | NZ_BALU01000025.1 |
| *L. crispatus* ST1 | NR_077026.1 |
| *L. gallinarum* DSM 10532<sup>T</sup> | NZ_AZEL01000029.1 |
| *L. helveticus* DPC 4571 | NR_076471.1 |
| PXN35 | PXN35 |

**Table 5.** PXN37 23S rRNA gene sequences used for the study.

| Strain | Sequence ID |
| --- | --- |
| PXN37 | PXN37 |
| *Lactcaseibacillus absianus* YH-lac23 | NZ_JABEKX010000006.1 |
| *Lactcaseibacillus chiayiensis* BCRC81062 | NZ_NOXN01000109.1 |
| *Lactcaseibacillus chiayiensis* NCYUAS | NZ_MSSM01000072.1 |
| *Lactcaseibacillus brantae* DSM23927<sup>T</sup> | NZ_AYZQ01000009.1 |
| *Lactcaseibacillus casei* ATCC393<sup>T</sup> | NZ_AP012544.1 |
| *Lactcaseibacillus casei* BIO5773 | NZ_WBOC01000014.1 |
| *Lactobacillus crispatus* ATCC33820<sup>T</sup> | NZ_AZCW01000100.1 |
| *Lactcaseibacillus paracasei* LOCK919 | NC_021721.1 |
| *Lactcaseibacillus paracasei* subsp *paracasei* JCM8130<sup>T</sup> | NZ_AP012541.1 |
| *Lactcaseibacillus rhamnosus* NCTC13764<sup>T</sup> | NZ_LR134331.1 |
| *Lactcaseibacillus rhamnosus* NRRL B-442 | NZ_PKQF01000030.1 |
| *Lactcaseibacillus saniviri* CJNU3004 | NZ_JAIZVU010000042.1 |
| *Lactcaseibacillus saniviri* DSM24301<sup>T</sup> | NZ_JQCE01000031.1 |

## RESULTS AND DISCUSSION

### Strain PXN35

Strain PXN35 was originally designated as *L. acidophilus* and deposited at National Collection of Industrial, Food and Marine Bacteria (NCIMB, Aberdeen, United Kingdom) as NCIMB 30184.

Due to *in vitro* microbiological and molecular evaluation discrepancies with *L. acidophilus* designation, almost full 16s rRNA gene sequencing was applied with a sample from the revived original master cell bank. Initially, *on line* tool BLAST grouped the strain with *Lactobacillus helveticus* sequences (data not shown), as a preliminary tentative identification. In order to confirm BLAST identification, a phylogenetic tree was constructed based on 16S rRNA with its closest phylogenetic neighbors. For the analysis, we selected the closest members of *Lactobacillus* phylogenetic group, using the most recent reclassification available and scientifically accepted [7]. The analysis grouped PXN35 sequence with all the *L. helveticus* sequences studied, including the species type strain sequence [7]. *Lactobacillus acidophilus* grouped separately. PXN35 and *L. helveticus* shared 99.43% 16s rRNA gene on average, whereas strain PXN35 only showed 97.83% homology with *L. acidophilus* representatives (Figure 1, Table 6).

**Figure 1.**
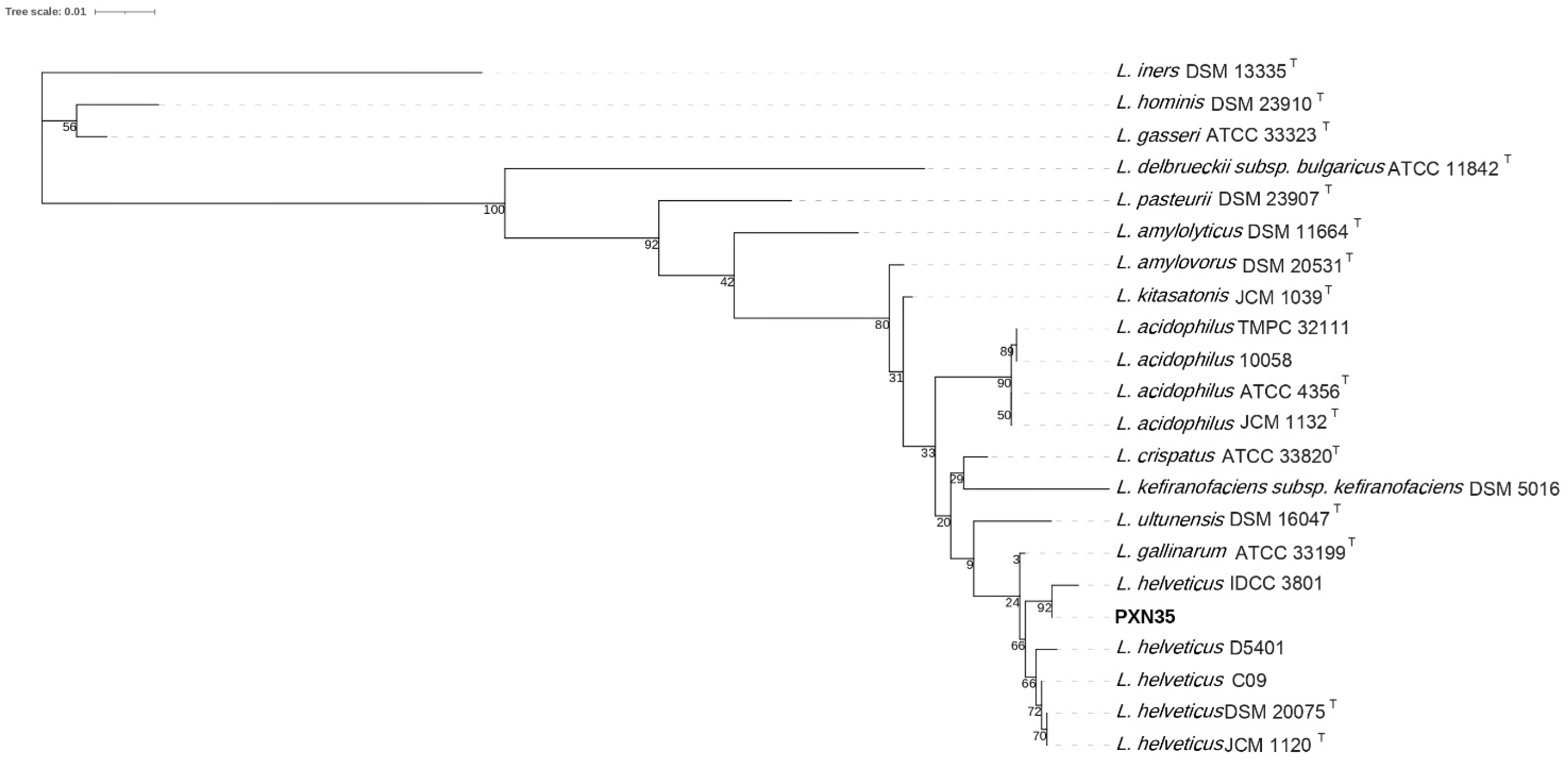
Phylogenetic tree constructed using the neighbour-joining method based on 16S rRNA gene sequences. Strain PXN35 sequence is close to *L. helveticus*. Bar, 10% nucleotide substitution.

**Table 6.** PXN35 16s rRNA gene % homology.

|  | *L. iners* strain DSM 13335 | *L. hominis* DSM 23910 | *L. gasseri* ATCC 33323 | *L. delbrueckii* subsp *bulgaricus* ATCC 11842 | *L. kefirano**faciens* subsp *kefirano**faciens* DSM 5016 | *L. acidophilus* strain TMPC 32111 | *L. acidophilus* strain 10058 | *L. acidophilus* ATCC 4356 | *L. crispatus* strain ATCC 33820 | *L. amylovorus* strain DSM 20531 | *L. kitasatonis* | *L. ultunensis* | *L. gallinarum* strain ATCC 33199 | *L. helveticus* DSM 20075 | *L. helveticus* strain C09 | *L. pasteurii* DSM 23907 | *L. amyolyticus* DSM 11664 | PXN35 |
| --- | --- | --- | --- | --- | --- | --- | --- | --- | --- | --- | --- | --- | --- | --- | --- | --- | --- | --- |
| *L. iners* strain DSM 13335 | 100.00 |  |  |  |  |  |  |  |  |  |  |  |  |  |  |  |  |  |
| *L. hominis* DSM 23910 | 94.50 | 100.00 |  |  |  |  |  |  |  |  |  |  |  |  |  |  |  |  |
| *L. gasseri* ATCC 33323 | 95.00 | 98.38 | 100.00 |  |  |  |  |  |  |  |  |  |  |  |  |  |  |  |
| *L. delbrueckii* subsp *bulgaricus* ATCC 11842 | 89.29 | 90.77 | 90.85 | 100.00 |  |  |  |  |  |  |  |  |  |  |  |  |  |  |
| *L. kefirano**faciens* subsp *kefirano**faciens* DSM 5016 | 89.52 | 91.00 | 90.93 | 92.06 | 100.00 |  |  |  |  |  |  |  |  |  |  |  |  |  |
| *L. acidophilus* strain TMPC 32111 | 89.57 | 91.27 | 91.13 | 92.33 | 96.60 | 100.00 |  |  |  |  |  |  |  |  |  |  |  |  |
| *L. acidophilus* strain 10058 | 89.87 | 91.70 | 91.57 | 93.05 | 97.24 | 99.29 | 100.00 |  |  |  |  |  |  |  |  |  |  |  |
| *L. acidophilus* ATCC 4356 | 89.94 | 91.78 | 91.64 | 93.13 | 97.31 | 99.22 | 99.93 | 100.00 |  |  |  |  |  |  |  |  |  |  |
| *L. crispatus* strain ATCC 33820 | 90.16 | 91.92 | 91.78 | 92.98 | 97.88 | 97.66 | 98.30 | 98.37 | 100.00 |  |  |  |  |  |  |  |  |  |
| *L. amylovorus* strain DSM 20531 | 89.66 | 91.71 | 91.57 | 92.70 | 97.45 | 97.52 | 98.16 | 98.23 | 98.73 | 100.00 |  |  |  |  |  |  |  |  |
| *L. kitasatonis* | 89.52 | 91.71 | 91.57 | 92.77 | 97.66 | 97.87 | 98.51 | 98.58 | 98.80 | 99.50 | 100.00 |  |  |  |  |  |  |  |
| *L. ultunensis* | 90.30 | 91.99 | 91.86 | 92.63 | 97.31 | 97.09 | 97.73 | 97.81 | 98.30 | 98.23 | 98.23 | 100.00 |  |  |  |  |  |  |
| *L. helveticus* strain IDCC 3801 | 89.57 | 91.27 | 91.21 | 91.70 | 95.96 | 97.30 | 97.02 | 96.95 | 97.09 | 96.88 | 96.88 | 96.95 |  |  |  |  |  |  |
| *L. gallinarum* strain ATCC 33199 | 90.30 | 92.20 | 92.07 | 93.13 | 97.31 | 97.73 | 98.37 | 98.44 | 98.58 | 98.23 | 98.37 | 98.44 | 100.00 |  |  |  |  |  |
| *L. helveticus* DSM 20075 | 90.16 | 92.06 | 91.93 | 92.84 | 97.03 | 97.59 | 98.23 | 98.30 | 98.30 | 97.95 | 98.09 | 98.16 | 99.58 | 100.00 |  |  |  |  |
| *L. helveticus* strain C09 | 90.23 | 92.13 | 92.00 | 92.77 | 97.10 | 97.66 | 98.30 | 98.37 | 98.37 | 98.02 | 98.16 | 98.23 | 99.65 | 99.93 | 100.00 |  |  |  |
| *L. pasteurii* DSM 23907 | 91.80 | 93.07 | 93.30 | 93.91 | 96.09 | 96.17 | 96.17 | 96.17 | 95.86 | 96.32 | 96.24 | 96.17 | 96.09 | 95.79 | 95.86 | 100.00 |  |  |
| *L. amylolyticus* DSM 11664 | 90.15 | 91.91 | 91.85 | 92.48 | 95.54 | 95.25 | 95.89 | 95.96 | 96.03 | 96.60 | 96.60 | 95.54 | 95.82 | 95.54 | 95.61 | 96.84 | 100.00 |  |
| PXN35 | 90.16 | 92.06 | 91.93 | 92.70 | 97.10 | 97.38 | 98.02 | 98.09 | 98.23 | 98.02 | 98.02 | 98.09 | 99.50 | 99.36 | 99.43 | 95.79 | 95.68 | 100.00 |

After that, we ran a further phylogenetic analysis with the same group of strains by comparing the almost full-length large ribosomal subunit 23S. Both 16S and 23S phylogenies are described to be in good agreement and provide an advantage over short ones like 5S, as local non-random rearrangements do not disturb the overall result [20]. The 23S phylogenetic tree structure closely resembled the 16S, with PXN35 grouped together with *L. helveticus* representatives and separated from *L. acidophilus*. When percentage homologies were studied, the analysis showed the highest nucleotide sequence homology between PXN35 and *L. helveticus* 23S rRNA gene (99.61%), whereas strain PXN35 showed 97.26% homology with *L. acidophilus* representatives (Figure 2, Table 7).

**Figure 2.**
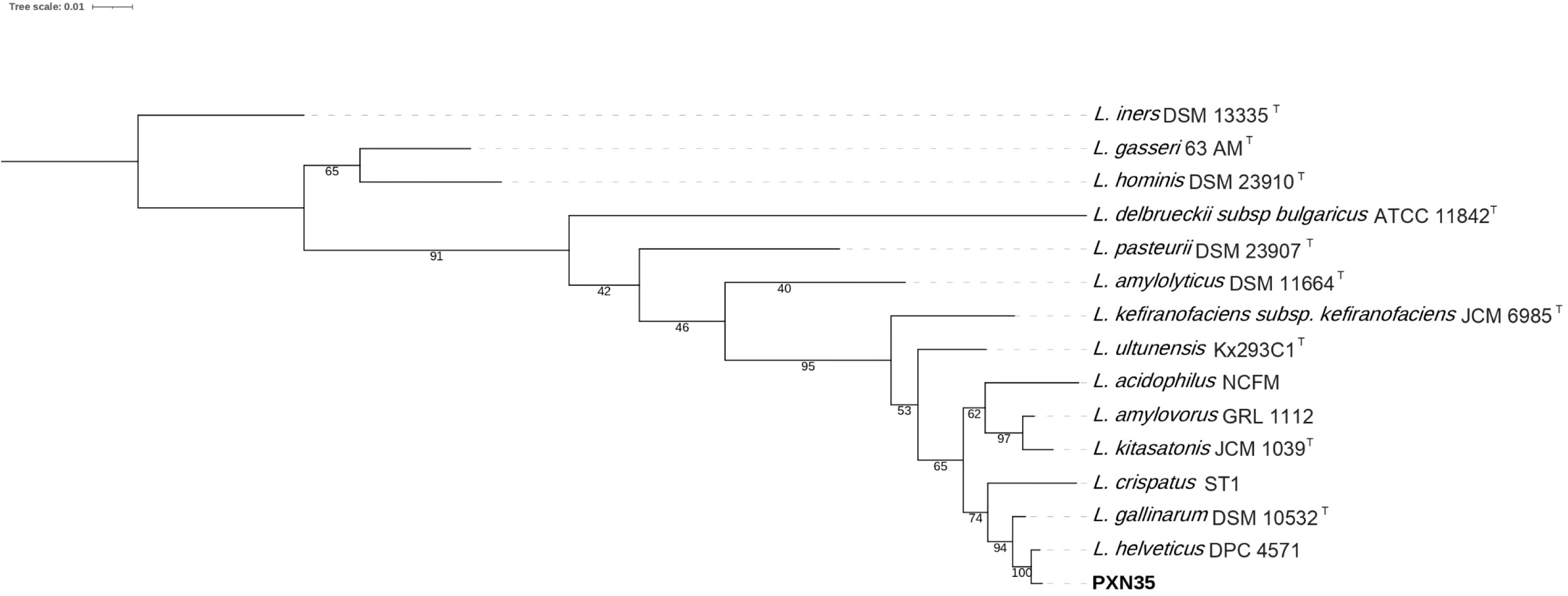
Phylogenetic tree constructed using the neighbour-joining method based on 23S rRNA gene sequences. Strain PXN35 sequence is close to *L. helveticus*. Bar, 10% nucleotide substitution.

**Table 7.** 23s rRNA gene % homology (only reference sequence is considered).

|  | *L. iners* DSM 13335 | *L. gasseri* 63 AM | *L. hominis* DSM 23910 | *L. delbrueckii* subsp *bulgaricus* strain ATCC 11842 | *L. amylolyticus* DSM 11664 | *L. pasteurii* DSM 23907 | *L. kefirano* subsp *kefirano* JCM 6985 | *L. ultunensis* Kx293C1 | *L. acidophilus* strain NCFM | *L. amylovorus* GRL 1112 | *L. kitasatonis* JCM1039 | *L. crispatus* ST1 | *L. gallinarum* DSM 10532 | *L. helveticus* DPC 4571 | PXN35 |
| --- | --- | --- | --- | --- | --- | --- | --- | --- | --- | --- | --- | --- | --- | --- | --- |
| *L. iners* DSM 13335 | 100.00 |  |  |  |  |  |  |  |  |  |  |  |  |  |  |
| *L. gasseri* 63 AM | 93.05 | 100.00 |  |  |  |  |  |  |  |  |  |  |  |  |  |
| *L. hominis* DSM 23910 | 92.27 | 95.65 | 100.00 |  |  |  |  |  |  |  |  |  |  |  |  |
| *L. delbrueckii* subsp *bulgaricus* ATCC 11842 | 89.47 | 88.97 | 89.24 | 100.00 |  |  |  |  |  |  |  |  |  |  |  |
| *L. amylolyticus* DSM 11664 | 89.24 | 90.15 | 90.07 | 89.49 | 100.00 |  |  |  |  |  |  |  |  |  |  |
| *L. pasteurii* DSM 23907 | 89.89 | 90.45 | 91.02 | 90.75 | 92.62 | 100.00 |  |  |  |  |  |  |  |  |  |
| *L. kefirano* subsp *kefirano* JCM 6985 | 89.46 | 90.33 | 90.47 | 89.24 | 92.98 | 92.12 | 100.00 |  |  |  |  |  |  |  |  |
| *L. ultunensis* Kx293C1 | 89.81 | 91.32 | 90.54 | 88.84 | 93.01 | 92.15 | 96.52 | 100.00 |  |  |  |  |  |  |  |
| *L. acidophilus* strain NCFM | 89.63 | 90.89 | 90.37 | 88.40 | 92.18 | 91.71 | 95.52 | 97.26 | 100.00 |  |  |  |  |  |  |
| *L. amylovorus* GRL 1112 | 89.81 | 91.29 | 90.47 | 89.15 | 92.84 | 92.64 | 96.00 | 96.87 | 97.43 | 100.00 |  |  |  |  |  |
| *L. kitasatonis* JCM1039 | 89.72 | 91.33 | 90.77 | 88.93 | 92.84 | 92.72 | 95.87 | 96.69 | 97.35 | 99.17 | 100.00 |  |  |  |  |
| *L. crispatus* ST1 | 89.72 | 90.76 | 90.51 | 89.15 | 92.89 | 92.29 | 95.78 | 96.30 | 96.69 | 97.00 | 97.13 | 100.00 |  |  |  |
| *L. gallinarum* DSM 10532 | 90.07 | 91.10 | 90.32 | 89.31 | 92.66 | 92.19 | 95.99 | 97.17 | 97.21 | 97.82 | 97.47 | 97.74 | 100.00 |  |  |
| *L. helveticus* DPC 4571 | 90.07 | 91.27 | 90.58 | 89.44 | 92.66 | 92.23 | 95.86 | 96.95 | 97.26 | 97.65 | 97.26 | 97.43 | 99.22 | 100.00 |  |
| PXN35 | 90.11 | 91.22 | 90.58 | 89.52 | 92.48 | 92.14 | 95.95 | 96.91 | 97.26 | 97.69 | 97.26 | 97.43 | 99.17 | 99.61 | 100.00 |

Isolated from a human individual and deposited in NCIMB collection as NCIMB 30184, strain PXN35 is a gram-positive, catalase negative strain. Cells are nonmotile and non-spore-forming, rod-shaped, isolated or tend to form pairs, with 1.0 μm x 5-15 μm. Strain is obligately homofermentative and grows well in commercially available media for lactic acid bacteria (MRS, Mann Rogosa and Sharpe) at 37°C. Colonies of strain PXN35 grown on MRS agar plates are grey, smooth, slightly domed, and round. The strain PXN35 has been shown to inhibit growth of *Salmonella Typhimurium, Clostridium difficile, Staphylococcus aureus, Escherichia coli* and *Enterococcus faecalis*. Strain PXN35 produces organic acids and significantly reduces pH [21, 22]. Originally designated as *L. acidophilus*, the present study demonstrates its unequivocal identification as *L. helveticus*.

### Strain PXN37

Originally identified as *L. casei*, strain PXN37 was deposited at NCIMB collection as NCIMB 30185. Further molecular evaluation with almost full 16s rRNA gene sequencing with the revived original master cell bank showed some divergences with *L. casei* species and BLAST tool tentatively grouped the strain with *Lacticaseibacillus paracasei* sequences (data not shown). The initial BLAST identification was checked by a phylogenetic tree based on 16S rRNA with is closest phylogenetic neighbors as Zheng and coworkers established [7]. Strain PXN37 was clearly grouped with all the *L. paracasei* sequences including the species type strain sequence [7] sharing 100% 16s rRNA gene on average with *L. paracasei* whereas *L. casei* representatives grouped separately (Figure 3, Table 8).**Rodenes et al. 2022**

**Figure 3.**
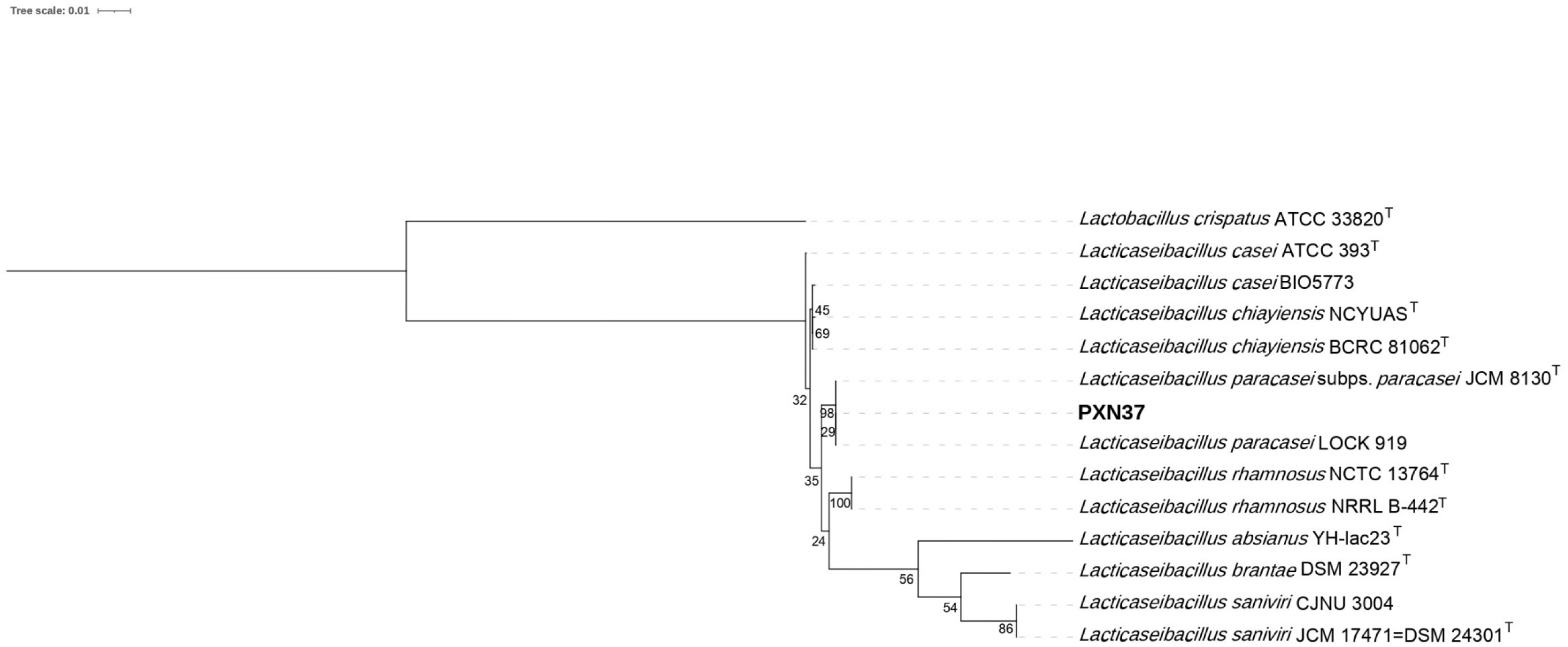
Phylogenetic tree constructed using the neighbour-joining method based on 16S rRNA gene sequences. Strain PXN37 sequence is close to *L. paracasei*. Bar, 10% nucleotide substitution.

**Table 8.** PXN37 16s rRNA gene % homology.

|  | *L. crispatus* ATCC33820<sup>T</sup> | *L. absianus* YH-lac23 | *L. rhamnosus* NCTC13764<sup>T</sup> | *L. rhamnosus* NRRL B-442 | *L. paracasei* subsp. *paracasei* JCM8130<sup>T</sup> | *L. paracasei* LOCK919 | *L. casei* ATCC393<sup>T</sup> | *L. chiayi*ensis NCYUAS | *L. chiayi*ensis BCRC81062 | *L. casei* BIO5773 | *L. saniviri* DSM 24301<sup>T</sup> | *L. saniviri* CJNU3004 | *L. brantae* DSM23927<sup>T</sup> | PXN37 |
| --- | --- | --- | --- | --- | --- | --- | --- | --- | --- | --- | --- | --- | --- | --- |
| *Lactobacillus crispatus* ATCC33820<sup>T</sup> | 100.00 |  |  |  |  |  |  |  |  |  |  |  |  |  |
| *Lacticaseibacillus absianus* YH-lac23 | 86.38 | 100.00 |  |  |  |  |  |  |  |  |  |  |  |  |
| *Lacticaseibacillus rhamnosus* NCTC13764<sup>T</sup> | 86.37 | 94.03 | 100.00 |  |  |  |  |  |  |  |  |  |  |  |
| *Lacticaseibacillus rhamnosus* NRRL B-442 | 86.37 | 94.03 | 100.00 | 100.00 |  |  |  |  |  |  |  |  |  |  |
| *Lacticaseibacillus paracasei* subsp. *paracasei* JCM8130<sup>T</sup> | 86.37 | 93.83 | 98.85 | 98.85 | 100.00 |  |  |  |  |  |  |  |  |  |
| *Lacticaseibacillus paracasei* LOCK919 | 86.37 | 93.83 | 98.85 | 98.85 | 100.00 | 100.00 |  |  |  |  |  |  |  |  |
| *Lacticaseibacillus casei* ATCC393<sup>T</sup> | 86.51 | 93.83 | 98.78 | 98.78 | 99.19 | 99.19 | 100.00 |  |  |  |  |  |  |  |
| *Lacticaseibacillus chiayi*ensis NCYUAS | 86.43 | 93.96 | 98.92 | 98.92 | 99.19 | 99.19 | 99.73 | 100.00 |  |  |  |  |  |  |
| *Lacticaseibacillus chiayi*ensis BCRC81062 | 86.43 | 93.89 | 98.92 | 98.92 | 99.19 | 99.19 | 99.73 | 99.93 | 100.00 |  |  |  |  |  |
| *Lacticaseibacillus casei* BIO5773 | 86.5 | 93.96 | 98.92 | 98.92 | 99.19 | 99.19 | 99.8 | 99.86 | 99.86 | 100.00 |  |  |  |  |
| *Lacticaseibacillus saniviri* DSM 24301<sup>T</sup> | 87.4 | 93.96 | 94.65 | 94.65 | 94.51 | 94.51 | 94.44 | 94.58 | 94.51 | 94.51 | 100.00 |  |  |  |
| *Lacticaseibacillus saniviri* CJNU3004 | 87.4 | 93.96 | 94.65 | 94.65 | 94.51 | 94.51 | 94.44 | 94.58 | 94.51 | 94.51 | 100.00 | 100.00 |  |  |
| *Lacticaseibacillus brantae* DSM23927<sup>T</sup> | 86.99 | 94.09 | 94.65 | 94.65 | 94.65 | 94.65 | 94.38 | 94.58 | 94.51 | 94.44 | 97.43 | 97.43 | 100.00 |  |
| PXN37 | 86.37 | 93.83 | 98.85 | 98.85 | 100.00 | 100.00 | 99.19 | 99.19 | 99.19 | 99.19 | 94.51 | 94.51 | 94.65 | 100.00 |

Further phylogenetic analysis with full-length large ribosomal subunit 23S was run as previously and as in the case of 16s rRNA analysis, PXN37 grouped together with *L. paracasei* representatives and separated from *L. casei*. When percentage homologies were studied, the analysis showed the highest nucleotide sequence homology between PXN37 and *L. paracasei* 23S rRNA gene (100%), and lower (98.59%) with *L. casei* representatives (Figure 4, Table 9).

**Figure 4.**
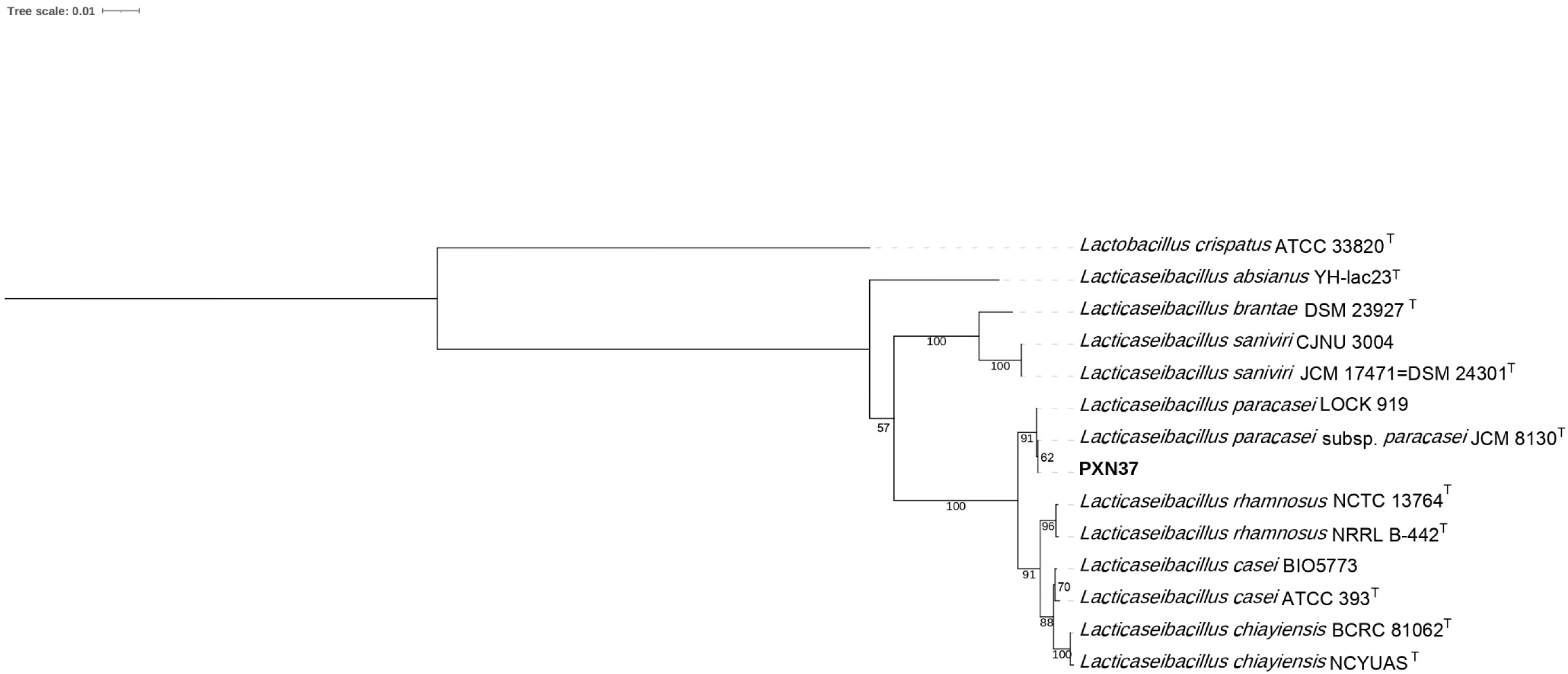
Phylogenetic tree constructed using the neighbour-joining method based on 23S rRNA gene sequences. Strain PXN37 sequence is close to *L. paracasei*. Bar, 10% nucleotide substitution.

**Table 9.** PXN37 23s rRNA gene % homology.

|  | *L. crispatus* ATCC33820<sup>T</sup> | *L. absianus* YH-lac23 | *L. paracasei* subsp *paracasei* JCM8130<sup>T</sup> | *L. paracasei* LOCK919 | *L. rhamnosus* NCTC13764<sup>T</sup> | *L. rhamnosus* NRRL B-442 | *L. casei* ATCC393<sup>T</sup> | *L. casei* BIO5773 | *L. chiayi*ensis BCRC81062 | *L. chiayi*ensis NCYUAS | *L. brantae* DSM23927<sup>T</sup> | *L. saniviri* CJNU3004 | *L. saniviri* DSM24301<sup>T</sup> | PXN37 |
| --- | --- | --- | --- | --- | --- | --- | --- | --- | --- | --- | --- | --- | --- | --- |
| *L. crispatus* ATCC33820<sup>T</sup> | 100.00 |  |  |  |  |  |  |  |  |  |  |  |  |  |
| *L. absianus* YH-lac23 | 83.85 | 100.00 |  |  |  |  |  |  |  |  |  |  |  |  |
| *L. paracasei* subsp *paracasei* JCM8130<sup>T</sup> | 83.85 | 93.48 | 100.00 |  |  |  |  |  |  |  |  |  |  |  |
| *L. paracasei* LOCK919 | 83.85 | 93.44 | 99.96 | 100.00 |  |  |  |  |  |  |  |  |  |  |
| *L. rhamnosus* NCTC13764<sup>T</sup> | 83.57 | 93.44 | 98.47 | 98.51 | 100.00 |  |  |  |  |  |  |  |  |  |
| *L. rhamnosus* NRRL B-442 | 83.65 | 93.48 | 98.51 | 98.55 | 99.88 | 100.00 |  |  |  |  |  |  |  |  |
| *L. casei* ATCC393<sup>T</sup> | 83.49 | 93.20 | 98.59 | 98.59 | 99.15 | 99.11 | 100.00 |  |  |  |  |  |  |  |
| *L. casei* BIO5773 | 83.49 | 93.24 | 98.59 | 98.59 | 99.15 | 99.11 | 99.84 | 100.00 |  |  |  |  |  |  |
| *L. chiayi*ensis BCRC81062 | 83.68 | 93.15 | 98.51 | 98.51 | 98.87 | 98.83 | 99.40 | 99.48 | 100.00 |  |  |  |  |  |
| *L. chiayi*ensis NCYUAS | 83.57 | 93.04 | 98.39 | 98.39 | 98.79 | 98.75 | 99.36 | 99.44 | 99.92 | 100.00 |  |  |  |  |
| *L. brantae* DSM23927<sup>T</sup> | 84.01 | 93.92 | 94.28 | 94.24 | 94.12 | 94.12 | 93.88 | 94.04 | 94.07 | 93.96 | 100.00 |  |  |  |
| *L. saniviri* CJNU3004 | 84.30 | 93.60 | 94.32 | 94.28 | 93.92 | 93.96 | 93.72 | 93.88 | 93.75 | 93.64 | 98.07 | 100.00 |  |  |
| *L. saniviri* DSM24301<sup>T</sup> | 84.30 | 93.60 | 94.32 | 94.28 | 93.92 | 93.96 | 93.72 | 93.88 | 93.75 | 93.64 | 98.07 | 100.00 | 100.00 |  |
| PXN37 | 83.85 | 93.48 | 100.00 | 99.96 | 98.47 | 98.51 | 98.59 | 98.59 | 98.51 | 98.39 | 94.28 | 94.32 | 94.32 | 100.00 |

The strain PXN37 was isolated from human origin as a gram-positive, catalase negative strain and deposited in NCIMB collection under reference NCIMB 30185. Cells are nonmotile and non-spore-forming, rod-shaped. Strain is obligately homofermentative and grows well in commercially available media for lactic acid bacteria (MRS, Mann Rogosa and Sharpe) at 37°C. The strain PXN37 has been shown to inhibit growth of *Salmonella Typhimurium, Clostridium difficile, Staphylococcus aureus, Escherichia coli* and *Enterococcus faecalis*. Strain PXN37 produces organic acids and significantly reduces pH [21,22]. Originally designated as *L. casei*, the present study demonstrates its unequivocal identification as *Lacticaseibacillus paracasei*.

## ACKNOWLEDGEMENTS

We thank Janice Rueda and Marta Tortajada for critical revision.

